# Transcription Factor (TF) validation using Dam-IT simultaneously captures genome-wide TF-DNA binding, direct gene regulation, and chromatin accessibility in plant cells

**DOI:** 10.1101/2025.05.06.652526

**Authors:** Will E. Hinckley, Austen Jack, Aurelia Li, Samantha Frangos, Angelo Pasquino, Shao-shan Carol Huang, Gloria M. Coruzzi

## Abstract

Transcription Factors (TFs) govern vast networks of gene regulation. However, TF-DNA binding and TF-gene regulation datasets are typically measured separately due to experimental constraints, making it challenging to disentangle true biological relationships from batch effects. To fill this gap, we developed <u>Dam</u>ID-seq <u>I</u>ncorporating <u>T</u>ranscriptomics (Dam-IT), which simultaneously captures TF-DNA binding, direct TF-gene regulation, and chromatin accessibility in the same batch of cells. Dam-IT uses a transient cell-based TF-target validation system that is scalable and flexible to many experimental designs. As proof of concept, we used Dam-IT to reveal that bZIP1 directly regulates genes by binding to DNA regions of relatively low chromatin accessibility, supporting a “Hit-and-Run” mechanism of transcription.

## Background

Comprehensive mapping of Transcription Factor (TF)-DNA binding is required to generate accurate gene regulatory networks (GRNs), but remains infeasible in many situations. DAP-seq is an *in vitro* method to map genome-wide direct TF-DNA binding sites in endogenous genome contexts.^1^ ChIP-seq captures information about *in vivo* cellular context, but requires antibodies, stable interactions, and more labor.^2^ TF Cut-&-Tag skips the immunoprecipitation, resulting in greater sensitivity compared to ChIP-seq while adhering to *in vivo* constraints. However, Cut-&-Tag still depends on antibodies and typically requires nuclei preparation.^3^

To address these limitations, DamID was developed to record a history of *in vivo* DNA binding for a given TF.^4^ DamID works by tethering a DNA Adenine Methyltransferase (DAM) to a TF and expressing this DAM-TF fusion *in vivo*. The DAM methylates adenines within GATCs near the sites of TF-DNA binding, resulting in a permanent DNA-binding footprint.^5^ DNA fragments containing adenine methylation can be specifically isolated and sequenced, resulting in genome-wide mapping of TF-DNA binding. Recently, single-cell Dam&T combined transcriptomics with single-cell DamID for chromatin protein profiling using a unique barcoding approach.^6^ They, along with others, show that DNA binding signal from an untethered DAM protein acts as a proxy for chromatin accessibility.^7^ Therefore, with proper controls, DamID can capture TF-DNA binding and chromatin accessibility *in vivo*.

We recently used DamID on the Arabidopsis TF *NLP7* to capture and map genome-wide transient TF-DNA binding.^8^ We now present an enhanced version, Dam-IT (DamID-seq Incorporating-Transcriptomics) which captures genome-wide TF-DNA binding, direct TF-gene expression regulation, and chromatin accessibility from the same batch of cells. Dam-IT greatly reduces genomic batch effects, and enables us to more accurately and rapidly characterize TF binding and regulation in the same sample of cells.

## Results and Discussion

Although DamID has been used to map protein-DNA interactions, broader adoptions are limited by high background and cellular toxicity resulting from persistent DAM methylation.^9^ To redress this, we previously developed a transient DamID system in which DAM-GR-TF fusion protein is transiently expressed in isolated plant cells.^10^ Dexamethasone (DEX) treatment allows for precise control of DAM-GR-TF nuclear import. Thus, adenine methylation at GATC sites is incorporated into the genome of the plant cells for only a brief time-period.

Adenine methylation by DAM-TF fusion proteins is a permanent footprint of TF-DNA binding. This contrasts with the binding signal of other TF-DNA binding assays that rely on immunoprecipitation. Methods such as ChIP-seq require time for chemical fixation, which biases results towards stable TF-DNA binding and misses transient TF-DNA interactions. Methylation footprinting by DamID technology results in greater detection of TF-DNA binding, as both stable and transient TF-DNA binding can be captured. The cumulative methylation signal also encompasses even the earliest TF-target gene interactions after DEX-mediated GR-TF nuclear import, which ChIP-seq would typically miss unless it was completed on fine time scales. We previously used this transient DamID approach to map the genome-wide binding of NLP7 in regulating early nitrogen signals in plant cells.^8^ We detected novel early and transient TF-DNA binding of NLP7-target genes that were missed by ChIP-seq. Importantly, DNA methylation by DamID captures a comprehensive *history* of both stable and transient TF-DNA binding in cells.

Our transient DamID approach parallels our published TARGET system (Transient Assay Reporting Genome-wide Effects of Transcription factors) that identifies direct TF-regulated genes.^11,12^ In the TARGET assay, mRNA profiles are measured following DEX-induction of GR-TF nuclear import, relative to empty vector (GR-EV) controls.^11,12^ Importantly, DEX-mediated GR-TF nuclear import occurs in the presence of cycloheximide (CHX). CHX inhibits the translation of TFs downstream of the perturbed TF. The combination of DEX and CHX treatments ensures detection of GR-TF directly-regulated target genes compared to GR-EV. We previously completed TARGET and DamID-seq on NLP7 to study its genome-wide “Hit-and-Run” activity.^8^ However, these experiments were not completed on the same cell samples. NLP7 DamID-seq and TARGET RNA-seq were compared from experiments completed on different days, meaning batch effects could confound our analyses. Our new Dam-IT assay enables us to measure TF-DNA binding and direct TF gene regulation in the same cell samples, as described below.

Our Dam-IT assay presented herein builds on our prior DamID-seq work^8^ with three key improvements: (1) incorporating Fluorescence Activated Cell Sorting (FACS) to isolate DAM-GR-TF transfected cells, (2) T7 exonuclease treatment of non-methylated DNA, and (3) collecting both the DNA fraction for TF binding signals and the mRNA fraction for gene expression from the same sample of cells. Adding the FACS step prevents DNA from untransfected cells from contaminating the final DNA library due to gDNA shearing. We found that incorporating FACS in DamID on NLP7 significantly increases the number of detected DNA binding peaks (Supp Table 1). Having established that FACS eliminates background from untransfected cells, we next aimed to reduce background within TF-transfected cells. We found this could be achieved by adding a T7 exonuclease treatment to degrade DNA fragments with no GATC methylation following the iDamID-seq protocol.^5^

The third update that completes our new Dam-IT method is the simultaneous collection of RNA and DNA fractions from the same cells, without the need for splitting cells into separate samples (Fig 1A). This is enabled by the Qiagen AllPrep Micro Kit (Qiagen 80284). Importantly, the DNA methylation signal from the DAM-GR-EV sample in combination with RNA from the same sample of cells also functions as a proxy of chromatin accessibility.^6^ Thus, Dam-IT captures genome-wide TF-DNA binding, direct TF-gene regulation, and chromatin accessibility from the same batch of cells. (Fig 1A)

**Figure 1:**
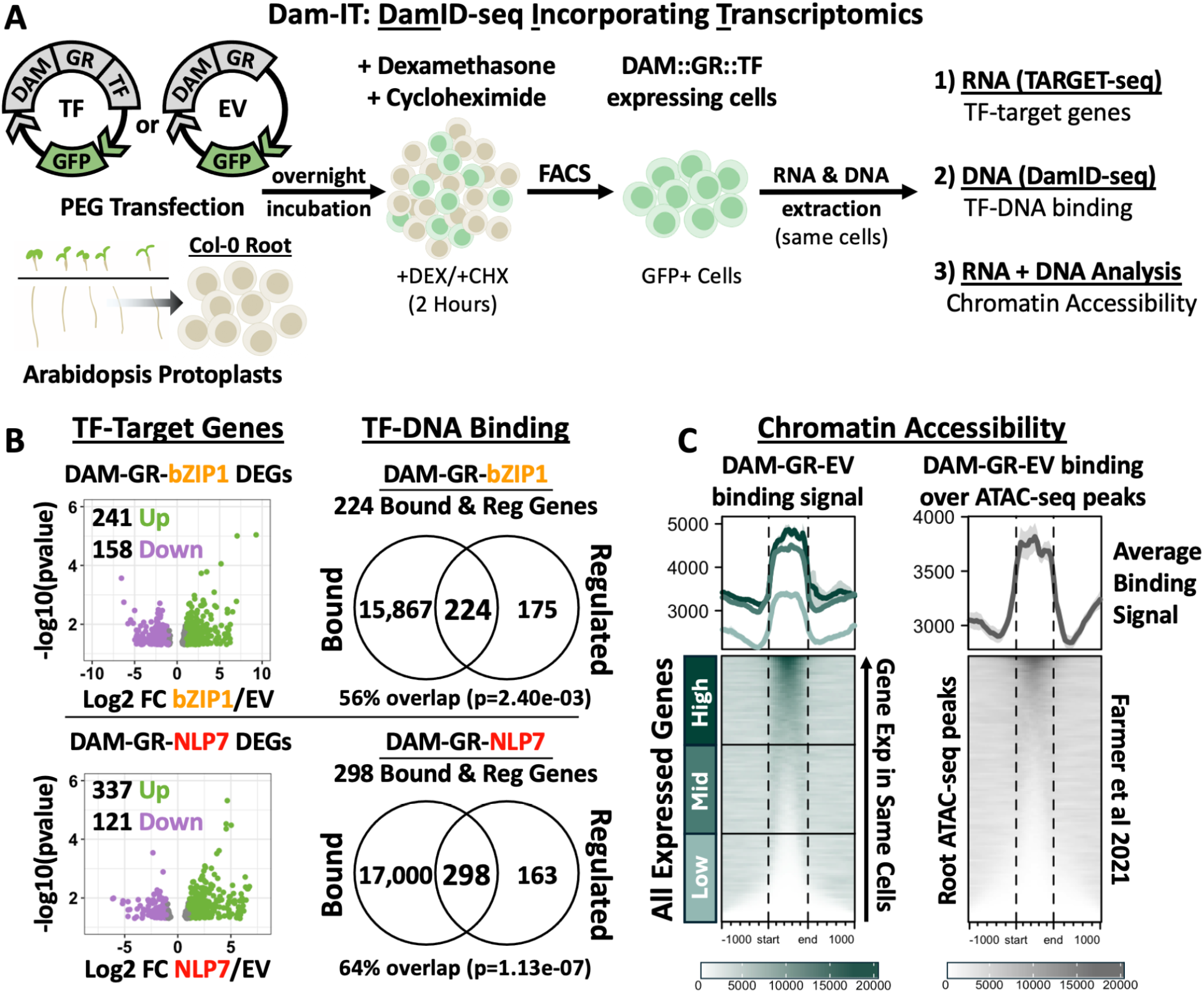
Dam-IT simultaneously captures TF-DNA binding, direct TF-gene regulation, and chromatin accessibility from the same batch of cells. **1A -** Dam-IT is a plant cell-based TF-perturbation method in which DAM:GR:TF fusion proteins are transiently expressed in plant protoplasts. DEX treatment triggers nuclear import of GR-TF fusion proteins, while simultaneous CHX treatment inhibits translation of secondary downstream TFs. Comparison to Empty Vector (DAM-GR-EV), allows the measurement of genome-wide direct gene regulatory activity of said TF.^11,12^ The Dam-IT protocol calls for FACS sorting of TF-transfected cells, and RNA and DNA are simultaneously extracted from the same sample of cells using the Qiagen AllPrep Micro kit. DNA and RNA fractions are then used for TARGET RNA-seq or DamID-seq as previously described (see Methods). **1B -** Dam-IT uncovered bZIP1 and NLP7 target genes that are TF-bound and directly TF-regulated in the same sample of cells. Genes are bound in at least one DamIT DNA binding replicate. Volcano plots show Log2FC DAM-GR-TF/DAM-GR-EV. Genes within 1,000 bp of DNA-binding peaks were intersected with TF-regulated genes. **1C -** The DAM-GR-EV (no TF) control from Dam-IT acts as a proxy for chromatin accessibility.^7^ Left: DAM-GR-EV DNA binding plotted over all expressed genes in the same cells, ordered by increasing gene expression. Right: DAM-GR-EV signal was plotted over Arabidopsis root ATAC-seq peaks from Farmer et al 2021, *Molecular Plant*.^14^

### Dam-IT TF-target direct regulation

We applied our Dam-IT method to two Arabidopsis nitrogen signal-regulating TFs, *bZIP1* and *NLP7*, each shown to bind transiently to early nitrogen-responsive “hit-and-run” targets.^8,13^ Root protoplasts were generated from Col-0 seedlings grown on low nitrogen media (1 mM KNO3). Approximately 10,000 transfected cells were collected per sample by FACS, and subjected to matched TARGET RNA-seq and DamID DNA-seq (Fig 1A, Supp Table 2). In the RNA-seq, each TF was found to evoke direct induction or repression of target genes relative to DAM-GR-EV controls. The Dam-IT method thus uncovered hundreds of directly-regulated DEGs for both bZIP1 and NLP7 (Fig 1B).

### Dam-IT TF-target DNA binding

DNA-binding peaks were uncovered for both TFs relative to DAM-GR-EV using a stringent q-value (q<1e08), and these peaks were located near thousands of Arabidopsis genes. This is expected, as TF-DNA binding is not broadly correlated to gene regulatory outcomes.^15^ Impressively, for both bZIP1 and NLP7, more than half of the directly regulated genes were also found to be TF-bound in the same sample of cells (Fig 1B). Previous ChIP-seq data for both TFs only found approximately 10% of TF-regulated genes to be TF-bound (ChIP-seq *in planta* for *NLP7*^*16*^, in cells for *bZIP1*^*13*^). In contrast to ChIP-studies, we note that Dam-IT uncovers 64% (NLP7) and 56% (bZIP1) of TF-regulated genes to be TF-bound in the same respective sample of cells.

### Dam-IT Chromatin Accessibility

We next examined the ability of Dam-IT to capture chromatin accessibility. As previously shown, increasing untethered DAM DNA-binding signal is positively correlated with chromatin accessibility and gene expression levels.^6^ DAM DNA-binding signal is correlated with ATAC-seq and FAIRE-seq, and DAM-DNA binding has been found to have greater resolution around less open chromatin.^7^ To test this in Dam-IT, we plotted the DNA methylation signal from DAM-GR-EV over all expressed genes, which were quantified in the same sample of cells. We ordered the genes by increasing gene expression levels, and interestingly, there is a clear positive correlation between DNA methylation in DAM-GR-EV samples and gene expression outcomes in the same sample of cells (Fig 1C). To further confirm this, we plotted DAM-GR-EV signal over ATAC-seq peaks from Arabidopsis roots, and found strong DAM-GR-EV methylation signal centered at the ATAC-seq peaks.^14^ We note that chromatin regions accessible in ATAC-seq but not in Dam-IT may be missed due to a lack of GATC sites, as GATC sites are required for the detection of methylation signals in Dam-IT.^7^ In sum, Dam-IT generates three important genomic data types; TF-DNA binding, direct TF-gene regulation and chromatin accessibility, all from the same batch of cells.

Our Dam-IT results shed new light on transient TF-target relationships that are not detectable by ChIP-seq. This was the first time DamID/Dam-IT was applied to *bZIP1*, a master nitrogen regulator that regulates genes via a “Hit-and-Run” mechanism.^13^ To look further into the genome-wide activity of *bZIP1*, we first analyzed bZIP1 DNA binding sequence preferences at Dam-IT bZIP1 binding sites. We queried the Dam-IT DNA binding peaks with the bZIP1 motif identified *in vitro* using DAP-seq.^1,2^ Using MEME suite tools^17^, we found significant enrichment of the bZIP1 binding motif among the bZIP1 Dam-IT DNA binding peaks (Fig 2A). This supports that bZIP1 DNA-binding detected by Dam-IT is associated with the canonical bZIP1 DNA-binding site.

**Figure 2:**
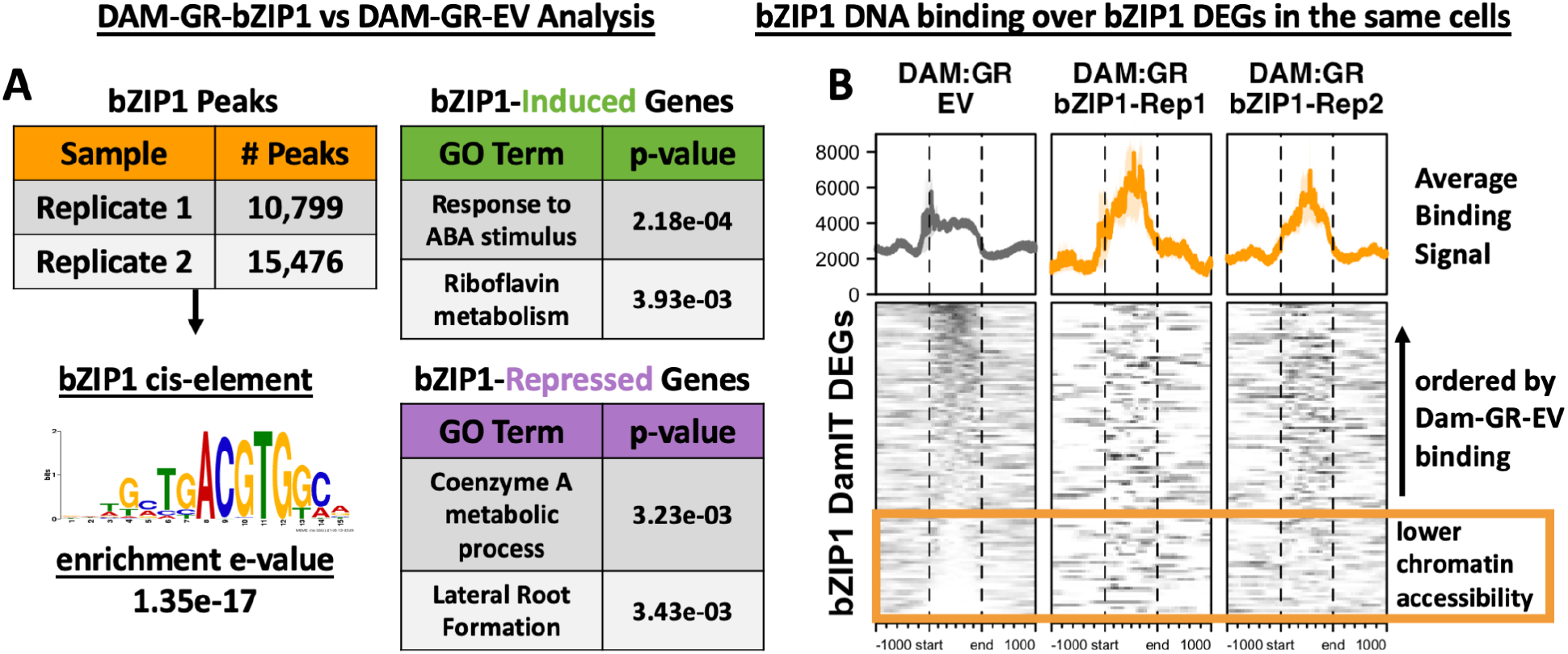
Dam-IT uncovers bZIP1 binding sites in regions of low chromatin accessibility associated with bZIP1 directly regulated genes. **2A -** bZIP1 DNA-binding peaks (q<1e-08) are significantly enriched for the bZIP1 DNA-binding motif. The e-value was calculated using SEA from MEME Suite Tools^17^ on the union of all bZIP1 peaks. Gene Ontology enrichment was calculated for the bZIP1 induced or repressed DEGs. **2B -** bZIP1 DEGs were ordered by DAM-GR-EV binding enrichment, where genes at the top of the heatmap have the highest DAM-GR-EV signal. This analysis found regions of DNA not accessible to DAM-GR-EV alone that were bound by DAM-GR-bZIP1 (gold box).

In our previous studies of bZIP1, we found that the majority of directly-regulated bZIP1 target genes were not detectably bound by ChIP-seq, likely due to transient binding.^13^ We hypothesized that this is due to the transient nature of bZIP1 DNA-binding in a “Hit-and-Run” mechanism of transcription.^18^ In the “Hit-and-Run” model of transcription, a TF binds to target genes just long enough to recruit transcriptional machinery before dissociating from DNA to regulate a different target gene.^19^ However, rapid and transient DNA-binding is difficult to detect genome-wide with ChIP-seq, as the assay steps require stable TF-target binding. By contrast, Dam-IT is capable of detecting such transient TF-DNA binding, as the adenine methylation is a permanent footprint of DNA binding. To test the bZIP1 “Hit-and-Run” hypothesis, we plotted bZIP1 Dam-IT DNA binding over the directly regulated bZIP1 target genes that were not bound by bZIP1 in ChIP-seq.^13^ We found evidence for enhanced Dam-IT bZIP1 binding to bZIP1 DEGs missed by ChIP-seq (Class III, Supp Figure 1). Thus, the Dam-IT approach enabled the discovery of bZIP1 DNA binding to bZIP1 DEGs that were missed by ChIP-seq, lending further support to the “Hit-and-Run” model of bZIP1 transient target binding and regulation.

Lastly, *bZIP1* is an early TF regulator of plant nitrogen responses.^20^ Our Dam-IT results indeed support the hypothesis that bZIP1 functions in a highly transient nature to rapidly initiate plant responses to nitrogen signals. We next explored if bZIP1 activity is constrained by chromatin inaccessibility, which could interfere with the rapid propagation of transcriptional responses to nitrogen. To test this, we plotted DAM-GR-bZIP1 DNA binding at the bZIP1 directly regulated genes (Dam-IT DEGs), ordered by DAM-GR-EV DNA binding signal enrichment as a proxy for accessible chromatin. Interestingly, we found DAM-GR-bZIP1 DNA binding in regions of chromatin near bZIP1-regulated genes that were inaccessible to DAM-GR-EV alone (Fig 2C, Gold Box). This implies that bZIP1 is required for the DAM to access these chromatin regions. This is the first evidence supporting that bZIP1 displays transcriptional pioneering activity, as previously hypothesized.^21,22^ bZIP1 binding regions of inaccessible chromatin is consistent with the proposed “Hit-and-Run” nature of bZIP1 activity, and supports the hypothesis that bZIP1 may be a Pioneer TF in the early nitrogen signaling pathway.^21,22^

## Conclusions

All together, our data show that Dam-IT is a novel method for measuring TF DNA binding, direct TF-gene regulation, and chromatin accessibility all from the same batch of cells. Beyond detecting stable TF-DNA binding, Dam-IT also captures early and transient TF targets, resulting in more comprehensive and informative GRNs. We used Dam-IT to validate transient bZIP1 DNA binding to bZIP1 directly regulated genes that were previously not detectable by ChIP-seq, supporting the “Hit-and-Run” model of bZIP1 activity. Lastly, as Dam-IT reveals TF-DNA binding in different chromatin states, Dam-IT will be a useful tool for characterizing pioneer activity across a wide range of plant TF families.

## Methods

### Plant growth & protoplast isolation

Col-0 *Arabidopsis thaliana* seeds were sterilized with 10% bleach and grown on 2% agar plates following the protocol outlined by Alvarez et al.^10^ (Supp Table 1), Col-0 and nlp7-1 mutant (SALK_026134C)^8^ protoplasts were used. The roots were removed from the plates after 10 days of growth. Protoplasts were prepared from the roots using 1% cellulase solution. Protoplasts were stained with FDA and counted on a hemocytometer under a fluorescent microscope. Enough protoplasts were produced for 3,000,000 cells for each treatment replicate.^12.13^

### Dam-IT Assay

3,000,000 *Arabidopsis* protoplasts cells isolated from roots were aliquoted for each treatment replicate. Each aliquot of cells was transfected with 120 ng of plasmid vector DNA mediated by PEG. The same vectors from Alvarez et al were used, which are available at ABRC.org under (CD3-2938 - CD3-2941).^8^ The transfected protoplasts were transferred to a 24-well cell culture plate and wrapped in foil to prevent degradation of mCherry fluorescence and incubated on a shake table overnight at room temperature for production of the desired DAM:GR:TF fusion proteins. Each replicate was treated with 5 µL of 1 mM CHX after the incubation and then treated with 1.25 µL 1 mM DEX two hours prior to FACS sorting on a BDFACSAria machine. FACS sorting was performed into 350 µl Qiagen AllPrep Buffer RLT+ which lyses cells and stabilizes nucleic acids for processing and analysis.

DNA and RNA from the FACS sorted cells were separated with the Qiagen AllPrep micro kit following the cell suspension protocol verbatim. DamID was performed on the DNA from each replicate as described in Alvarez et al., except for the addition of the previously described FACS sorting and T7 exonuclease degradation of background DNA for all replicates (Alvarez et al., 2023).^10^ DNA libraries were prepared with the NEBNext Ultra II DNA library prep kit following manufacturer’s protocol as described. RNA libraries were prepared with the NEB Ultra II RNA Library Prep kit following manufacturer’s protocol. Both the DNA and RNA libraries were analyzed on Tapestation for the proper library fragment sizes. DNA and RNA libraries were sequenced with Illumina. Single-end sequencing was used for DNA, and paired-end sequencing was used for RNA. Library fragment length was set to 150 bp for DNA library sequencing and 100 bp for RNA library sequencing.^11.12^

### DNA Analysis

DNA library FASTQ files were first trimmed with CutAdapt with the default parameters as well as the following: “-q 30 --times 4 --trim-n -m 25”. The trimmed files were processed in FastQC with default parameters to ensure quality of trimmed sequences. Trimmed files were aligned to the TAIR10 Arabidopsis genome (Araport11 annotation) using HiSat2 with the default parameters, then sorted into BAM files in SamTools. DeepTools bamcompare was used to produce BigWig files with default parameters as well as the following: “--binSize 1 --minMappingQuality 30 --normalizeUsing RPKM -e 300”. These BigWig files were used to visualize peaks in JBrowse. Sorted BAM files were filtered for mapping quality in SamTools with SamTools view and the following parameter: “-q 30”. For each sample, the two replicates of filtered BAM files were merged with SamTools merge, which were used in downstream analysis. All filtered, merged, and intersect BAM files were processed in MACS2 to call peaks. MACS2 predictd was used to determine the “--extSize” parameter for each sample which will be used with MACS callpeak. MACS2 callpeak was used for each sample bam file relative to DAM-GR-EV with the following parameters: “-f BAM -g 1.2e8 --nomodel --extsize (predictd value) -q e-8”. BedTools intersect was used to find common peaks between the two filtered BAM files for each replicate. The first three columns of resulting narrowPeak files were processed in BedTools with the Araport11_protein_coding.201606.bed *Arabidopsis* gene annotation. BedTools window was run for each bed file with parameters “-sw -r 1000 -l 0” to intersect peaks with genes. The final gene lists were analyzed in R for peak visualization and intersection with RNA-seq data.

### RNA Analysis

RNA-seq library FASTQ files were trimmed with CutAdapt with the default parameters as well as the following: “-q 30 --trim-n -m 25”. Quality of trimmed RNA library reads was analyzed with FastQC at default parameters. The trimmed reads were aligned to the TAIR10 *Arabidopsis* genome with STAR paired-end alignment on default parameters. Subread FeatureCounts with the paired-end parameter was used to count how many reads were aligned to each gene in the *Arabidopsis* genome. Final gene counts files were analyzed in R with DESeq2 for differentially expressed genes (DEGs, p<0.05).The clusterProfiler package was used to gene ontology, and EnrichedHeatmap was used to plot DNA binding over the TF-regulated genes.

## Acknowledgments

This work was supported in part through the NYU IT High Performance Computing resources, services, and staff expertise. Sequencing and FACS sorting was performed by the NYU Genomics Core Facility with generous DNA sequencing subsidies from the Zegar Family Foundation.

## Funding

This work was supported by the NIH award R35GM138143 to S.C.H., NIH-NIGMS 1 R01 GM121753 to G.M.C, QBIST NIH T32 Training Program Grant 1T32GM132037-01 of NYU Biology awarded to W.E.H.

## Supplementary Figures and Tables

**Supplemental Figure 1:**
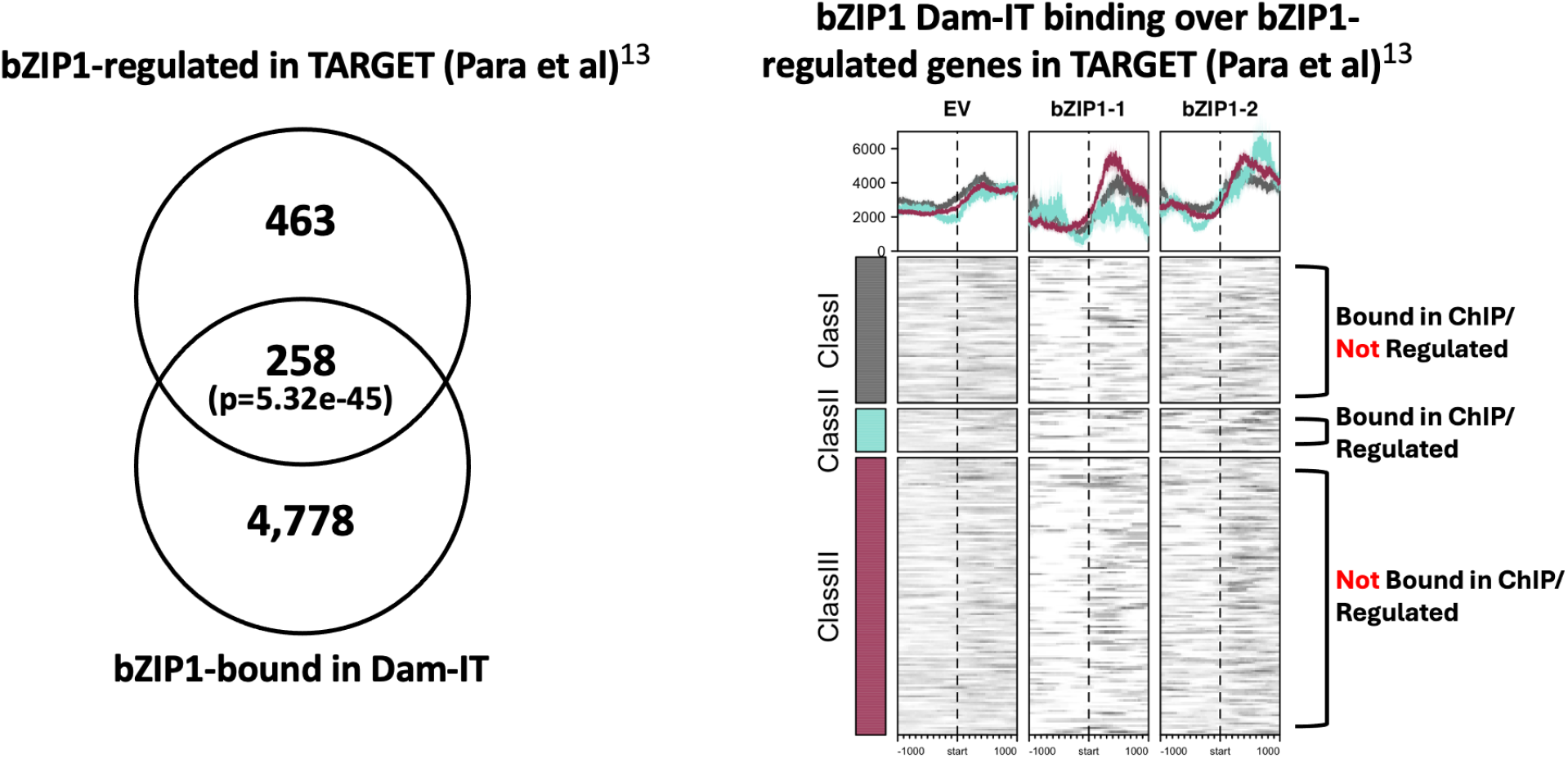
DAM-GR-bZIP1 binds bZIP1 direct regulated target genes that were previously missed by ChIP-seq (Para et al).^13^. **A)** bZIP1 Dam-IT bound genes significantly intersect with bZIP1 DEGs from Para et al 2014. **B)** DAM-GR-bZIP1 binds to ClassIII bZIP1 TARGET genes not bound by bZIP1 in ChIP-seq.

**Supplemental Table 1:** Dam-ID peaks called for DAM-GR-NLP7 binding in the presence/absence of FACS sorting of TF-transfected cells.

| Genotype | FACS | Sample | # Peaks (q<0.05) |
| --- | --- | --- | --- |
| *nlp7* mutant | Not Sorted | NNN1 | 376 |
|  |  | NNN2 | 339 |
|  |  | NNN3 | 579 |
| WT | Not Sorted | NNW1 | 1,042 |
|  |  | NNW2 | 404 |
|  |  | NNW3 | 721 |
| *nlp7* mutant | Sorted | SNN1 | 1,900 |
|  |  | SNN2 | 2,405 |
|  |  | SNN3 | 1,856 |
| WT | Sorted | SNW1 | 2,200 |
|  |  | SNW2 | 2,493 |
|  |  | SNW3 | 2,657 |

**Supplemental Table 2:** Dam-IT uncovers both TF-DNA binding and TF directly regulated DEGs from the same samples of cells.

**A) DNA Alignment Data**
| DNA Alignment Data | # Reads | % Reads Passing Filter | % Aligned Reads | # Peaks Called | Intersect Peaks |
| --- | --- | --- | --- | --- | --- |
| bZIP1-1 | 21,590,774 | 88.70% | 83.96% | 10,193 | 1798 |
| bZIP1-2 | 17,517,353 | 90.50% | 87.38% | 13,766 |  |
| bZIP1-merged | NA | NA | NA | 16,339 | NA |
| NLP7-1 | 12,511,330 | 92.80% | 88.72% | 10,799 | 1885 |
| NLP7-2 | 20,188,912 | 92.90% | 86.20% | 15,476 |  |
| NLP7-merged | NA | NA | NA | 16,378 | NA |
| Dam-GR-EV-1 | 19,081,274 | 93.60% | 90.25% | 21,537 | NA |

**B) RNA Alignment Data**
| RNA Alignment Data | # Raw Reads | % Passing Filters | # Aligned Reads | Overall Alignment Rate | # DEGs |
| --- | --- | --- | --- | --- | --- |
| bZIP1-1 | 21,397,001 | 63.80% | 7,589,549 | 55.58% | 399 |
| bZIP1-2 | 13,100,295 | 77.40% | 2,676,894 | 26.41% |  |
| NLP7-1 | 13,405,373 | 73.30% | 6,404,726 | 62.11% | 461 |
| NLP7-2 | 13,657,100 | 79.10% | 7,691,048 | 67.84% |  |
| Dam-GR-EV-1 | 18,116,984 | 77.30% | 9,339,993 | 66.71% | NA |
| Dam-GR-EV-2 | 10,049,407 | 73.50% | 5,276,667 | 68.57% |  |

## Notes

### Competing Interest Statement

The authors have declared no competing interest.

